# Nuclear Transport Factor 2 (NTF2) suppresses metastatic melanoma by modifying cell migration, metastasis, and gene expression

**DOI:** 10.1101/2020.03.13.991141

**Authors:** Lidija D. Vuković, Karen H. White, Jason P. Gigley, Daniel L. Levy

## Abstract

While changes in nuclear structure and organization are frequently observed in cancer cells, relatively little is known about how nuclear architecture impacts cancer progression and pathology. To begin to address this question, we studied Nuclear Transport Factor 2 (NTF2) because its levels decrease during melanoma progression. We show that increasing NTF2 expression in metastatic melanoma cells reduces cell proliferation and motility while increasing apoptosis. We also demonstrate that increasing NTF2 expression in these cells significantly inhibits metastasis and increases survival of mice. Mechanistically, we show that NTF2 levels affect the expression and nuclear positioning of a number of genes associated with cell proliferation and migration. We propose that by decreasing nuclear size and/or lamin A nuclear localization, ectopic expression of NTF2 in metastatic melanoma alters chromatin organization to generate a gene expression profile with characteristics of primary melanoma, concomitantly abrogating several phenotypes associated with advanced stage cancer both *in vitro* and *in vivo*. Thus NTF2 acts as a melanoma tumor suppressor to maintain proper nuclear structure and gene expression and could be a novel therapeutic target to improve health outcomes of melanoma patients.

## INTRODUCTION

Cancer cells exhibit many characteristics that distinguish them from normal cells, including higher proliferation rates, immortalization, suppressed apoptosis, and metastasis (Weinberg, 2014). In addition to these functional differences, cancer cells are also morphologically distinct, with irregularly shaped cells, increased nuclear-to-cytoplasmic ratios, prominent nucleoli, and larger nuclei. Pathologists have long used altered nuclear morphology to distinguish cancer cells from normal cells, and late stage tumors often exhibit more aberrant nuclear structures compared to earlier stages (Bignold et al., 2006; Chow et al., 2012; Jevtic and Levy, 2014; Zink et al., 2004). Nuclear size has diagnostic and prognostic value in melanoma (Mossbacher et al., 1996; Sorensen and Erlandsen, 1990; Stolz et al., 1987), one of the most prevalent cancers with cases of cutaneous melanoma growing at the fastest rate of all cancers among Caucasians (Leiter and Garbe, 2008). However, how altered nuclear architecture contributes to progression and pathology of a broad range of cancers, including melanoma, is unknown. The six major steps of melanoma progression are: common acquired nevus without cytogenetic abnormalities, melanocytic nevus with aberrant differentiation, dysplastic nevus, radial growth phase (RGP) primary melanoma, vertical growth phase (VGP) primary melanoma, and metastatic melanoma (Clark et al., 1984). In VGP melanoma some local dermal cell invasions are possible, while cell dissemination over longer distances is a defining characteristic of metastatic melanoma (Ciarletta et al., 2011).

We previously reported that RGP primary melanoma cells exhibit larger nuclei compared to normal melanocytes. We also found that the levels of Nuclear Transport Factor 2 (NTF2) inversely correlate with nuclear size, with decreasing NTF2 levels in melanoma progression correlating with increased nuclear size (Vukovic et al., 2016). NTF2 plays a critical role in nucleocytoplasmic transport by recycling RanGDP from the cytoplasm to the nucleus where GDP is exchanged for GTP by chromatin-bound RCC1 (Clarkson et al., 1996; Corbett and Silver, 1996; Madrid and Weis, 2006; Paschal and Gerace, 1995). NTF2 is also associated with the nuclear pore complex (NPC) where it limits cargo import based on cargo size (Bayliss et al., 1999; Feldherr et al., 1998). In particular NTF2 restricts import of nuclear lamins, intermediate filament proteins that line the inner nuclear membrane and regulate chromatin organization (Dittmer and Misteli, 2011). By reducing lamin import, NTF2 decreases nuclear growth and size (Levy and Heald, 2010). Based on transient transfections in different cancer and normal cell lines, we found that simply increasing NTF2 levels is sufficient to reduce nuclear size (Vukovic et al., 2016).

In this study, we investigated if increasing NTF2 expression can impact the pathogenesis of metastatic melanoma. To address this question, we used two melanoma cell lines isolated from the same patient corresponding to VGP primary melanoma (WM983A) and metastatic melanoma (WM983B). We generated a stably transfected version of the metastatic melanoma cell line in which we could titrate the NTF2 expression level. Our data demonstrate that increased NTF2 expression reduces cell motility and cell proliferation while increasing apoptosis. In mice injected with these melanoma cells, increasing NTF2 levels reduces lung metastases and prolongs survival. Furthermore, RNA sequencing of cells expressing different levels of NTF2 revealed differentially expressed genes involved in cell migration and cell survival. For a subset of these genes, we demonstrate that NTF2-dependent changes in gene positioning within the nuclear space correlate with changes in nuclear size and lamin A nuclear localization. Lastly, we show that VGP primary melanoma cells express higher NTF2 levels relative to metastatic melanoma and that increasing NTF2 expression in metastatic melanoma recapitulates some cellular phenotypes and gene expression patterns characteristic of VGP primary melanoma. We propose that NTF2 affects gene positioning through changes in lamin A nuclear localization and/or nuclear size, leading to altered expression of genes involved in cell proliferation, migration, and metastasis. These results highlight NTF2 as a potential novel therapeutic target to treat metastatic melanoma.

## RESULTS

### Increased NTF2 expression reduces cell proliferation and increases apoptosis

We previously published that expression of the nuclear transport protein NTF2 inversely correlates with nuclear size during melanoma progression (Vukovic et al., 2016). A critical unanswered question is if changes in nuclear architecture and nucleocytoplasmic transport have any relevance to cancer progression and disease severity. Therefore we developed stably transfected metastatic melanoma WM983B cells that allow for titratable expression of NTF2 dependent on doxycycline dosage. With 20 ng/ml doxycycline added to the growth media, on average 7.8-fold higher NTF2 protein expression was observed compared to the parental cell line (Fig. 1A). Interestingly, stably transfected cells not treated with doxycycline expressed ∼4-fold higher and more variable NTF2 protein levels compared to the parental cell line, an indication of promoter leakiness that has been reported by others (Cheng et al., 2015). For the sake of simplicity, we will refer to the parental cell line WM983B as “NTF2 low”, stably transfected WM983B without doxycycline as “NTF2 high dox-”, and stably transfected WM983B with 20 ng/ml doxycycline as “NTF2 high dox+” (Fig. 1A). To confirm promoter leakiness, we transiently transfected NTF2 high dox-cells with a commercially available pTet-tTS plasmid to suppress the leakiness, which reduced the NTF2 level to that of the parental cell line (data not shown).

**Figure 1:**
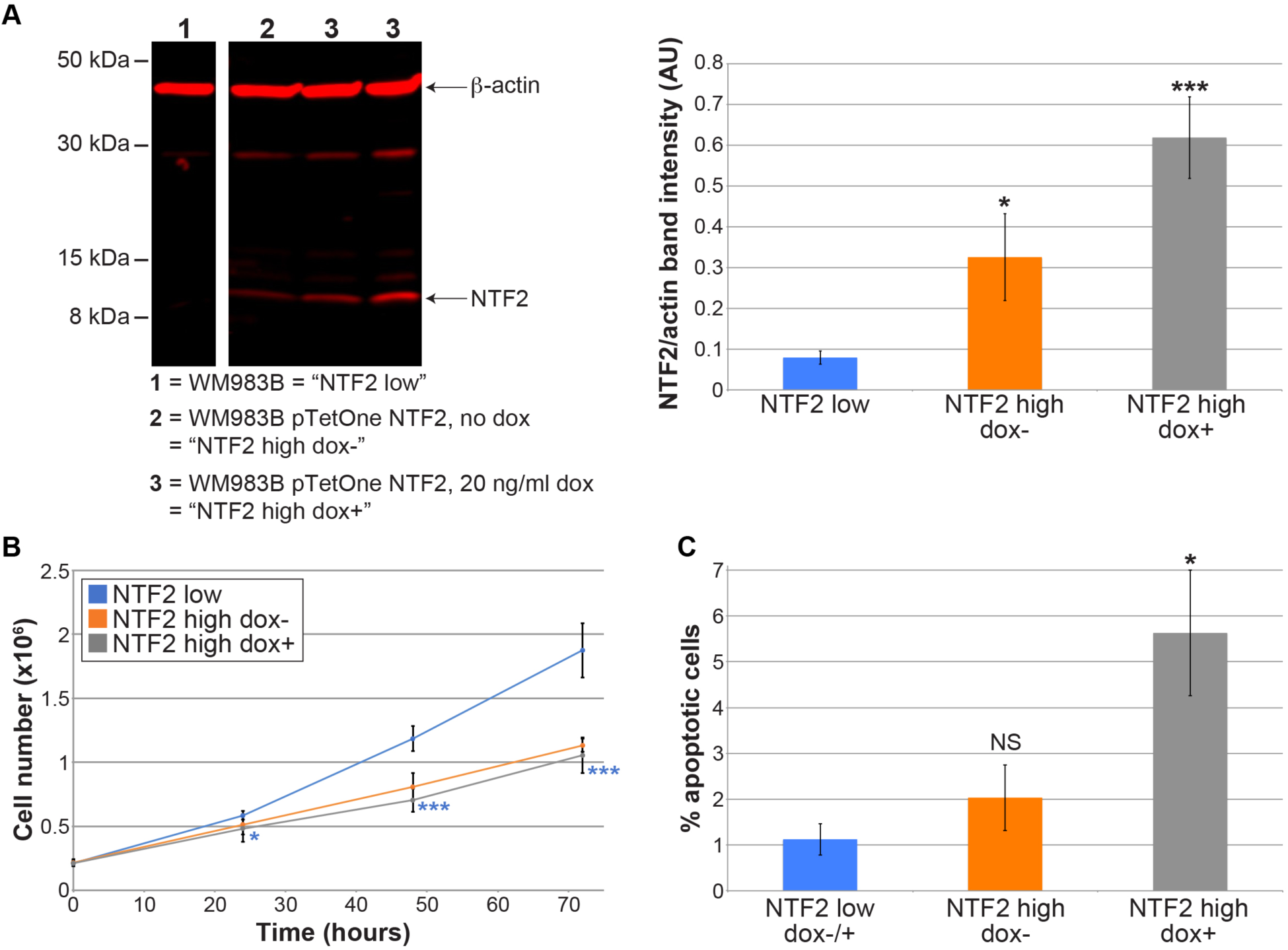
Inducing NTF2 expression in metastatic melanoma reduces cell proliferation and increases apoptosis. **(A)** Metastatic melanoma cells (WM983B) were stably transfected with a doxycycline-inducible NTF2 construct (pTetOne NTF2). These cells were treated with different concentrations of doxycycline for 48 hours and cell lysates were analyzed by NTF2 and actin immunoblots. For simplicity, “NTF2 low” refers to the parent cell line WM983B, “NTF2 high dox-“ refers to the transfected cell line grown without doxycycline, and “NTF2 high dox+” refers to the transfected cell line treated with 20 ng/µl doxycycline. NTF2 band intensities were normalized to actin band intensities and averaged for 7-8 independent samples per condition. Error bars are SEM. * p < 0.05, *** p < 0.005. The 20 ng/µl doxycycline concentration was selected for all experiments because it induces the greatest reduction in nuclear size. **(B)** 2×10^5^ cells were plated in 6-well plates. At 24, 48, and 72 hours, cells were trypsinized, stained with trypan blue, and counted. Data are from 3-4 independent experiments and two replicates per experiment. Error bars are SD. * p < 0.05, *** p < 0.005. The statistical comparisons are for NTF2 high dox-/+ compared to NTF2 low. **(C)** Cells were plated at 30% confluency. After 48 hours, cells were stained for activated caspase-3/7. The number of positive staining cells and total cells were counted to calculate the percent of apoptotic cells. For each condition, 8-16 coverslips were examined and 20-632 cells (140 cells on average) were counted per coverslip. Error bars are SEM. * p < 0.05, NS not significant.

We examined how increased NTF2 expression in metastatic melanoma affected various cancer cell characteristics. We first measured cell proliferation rates. Cells were plated at a constant density and then counted 24, 48, and 72 hours later. There were 44% fewer NTF2 high dox+ cells at 72 hours compared to NTF2 low cells (Fig. 1B). We next quantified the fraction of apoptotic cells by staining for activated caspase-3/7. Compared to NTF2 low cells, the apoptosis frequency was 5-fold higher in NTF2 high dox+ cells (Fig. 1C). To test if NTF2 has a similar effect in a different type of cancer, we increased NTF2 expression in LNCaP prostate cancer cells. Consistent with the melanoma results, ectopic NTF2 expression reduced cell proliferation in cultured prostate cancer cells (Fig. S1A-C). These data show that increasing NTF2 levels leads to decreased cell proliferation and increased apoptosis.

### Increased NTF2 levels reduce melanoma cell motility

Acquisition of metastatic potential by melanoma cells is the critical change that leads to patient death (Schadendorf et al., 2018). To initially test if NTF2 expression impacts metastasis, we performed *in vitro* would healing cell migration assays. We created small scratches in confluent monolayers of NTF2 low, NTF2 high dox-, and NTF2 high dox+ melanoma cells and observed cell behavior by time-lapse microscopy. The NTF2 low cell line was highly motile, completely filling the scratch after only 3 hours (Fig. 2A). During that same 3-hour time period, the NTF2 high dox- and NTF2 high dox+ cells only covered 43% and 11% of the scratch area, respectively. We repeated these experiments with a larger scratch area and increased the observation time to 10 hours. The same trend was observed with the NTF2 low, NTF2 high dox-, and NTF2 high dox+ cells filling 70%, 38%, and 21% of the scratch area, respectively (Fig. 2B and Movies 1-2). These data reveal an inverse correlation between NTF2 expression levels and cell migration potential. In these experiments we did not inhibit cell division since even low concentrations of drugs that affect microtubules can impact cell migration (Yang et al., 2010). Because filling of the scratch in our wound healing assays might be affected by cell proliferation, we also examined migration of non-dividing individual cells on the edge of the scratch (Fig. 2C). The NTF2 low cell trajectories were the longest with the NTF2 high dox- and NTF2 high dox+ cells exhibiting progressively reduced cell movements (Fig. 2D). These data show that titrated increases in NTF2 levels concomitantly decrease cell motility.

**Figure 2:**
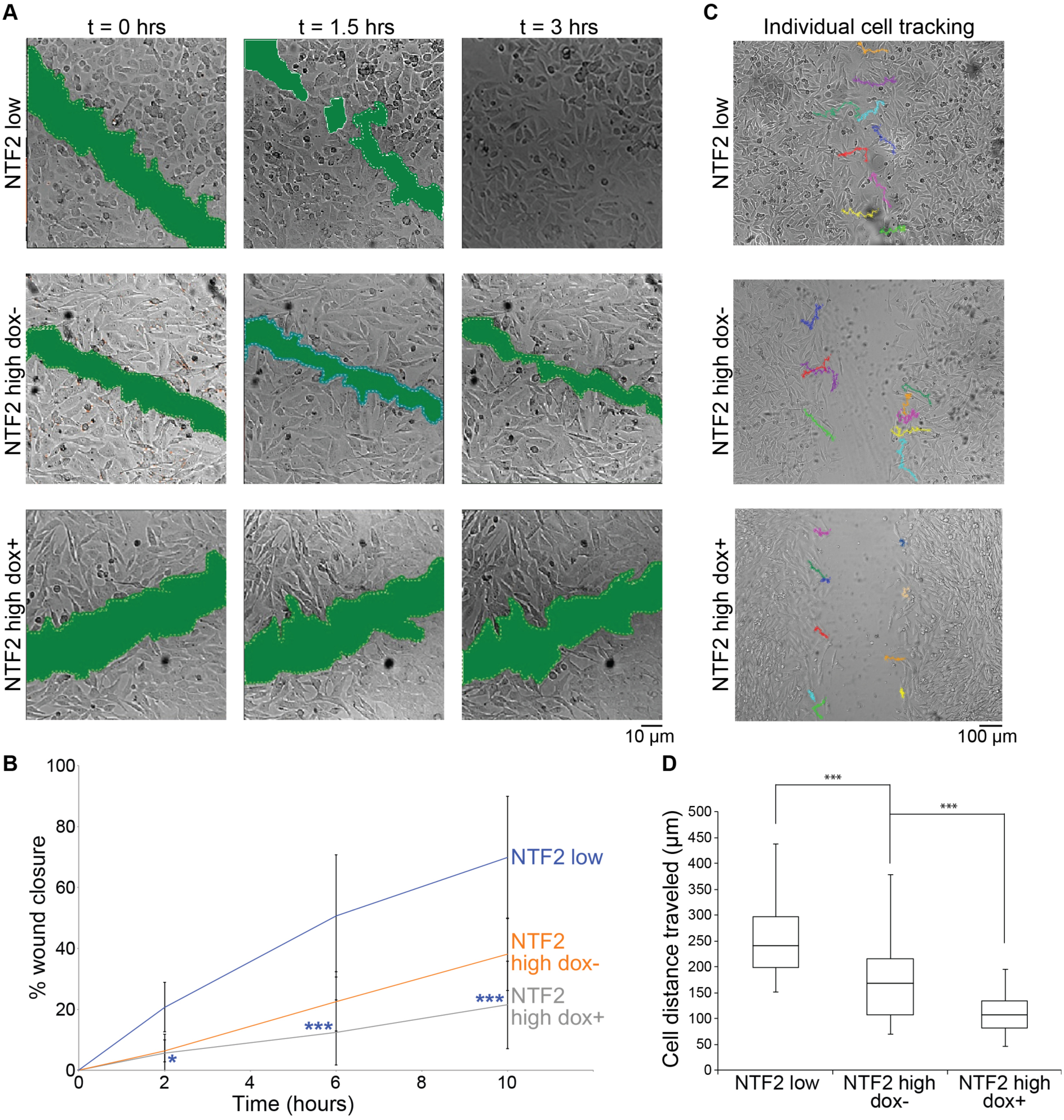
Higher levels of NTF2 suppress melanoma cell motility. **(A)** Wound healing assays were performed by creating small scratches (i.e. P2 pipet tip) through completely confluent cell monolayers and imaging over 3 hours. Green patches generated in Metamorph show the scratch area size. **(B)** Wound healing assays were performed by creating large scratches (i.e. P20 pipet tip) through completely confluent cell monolayers and imaging at 2, 6, and 10 hours. The percent wound closure was calculated relative to the scratch area at t=0. For each cell line, 5-8 independent experiments were performed. P-value asterisks indicate statistical differences between NTF2 low and NTF2 high dox+ cells. **(C)** The migration trajectories of individual cells in wound healing assays were measured over 10 hours using ImageJ. **(D)** The experiments described in (C) were quantified by measuring the total distance traveled by individual cells. For each condition, 44-57 cells were tracked. Error bars are SD. *** p < 0.0001; * p < 0.005.

### Increased NTF2 levels reduce melanoma metastasis *in vivo*

Since we observed that NTF2 expression levels influence *in vitro* cell migration, we next examined the metastatic potential of these cell lines *in vivo*. For these experiments, 6-8 week old Rag2^−/−^ γc^−/−^ knockout mice (i.e. deficient for B, T, and innate lymphoid cells) were intravenously injected with NTF2 low or NTF2 high cells. The latter group was divided into two cohorts, in which one was provided with doxycycline-containing food from the first day of injection and labeled as NTF2 high dox+. The group not treated with doxycycline was labeled NTF2 high dox-. Four months post-injection, the mice were sacrificed and lungs and other internal organs were examined for tumors. In the NTF2 low and NTF2 high dox-mice, tumors were always found on the lungs (Fig. 3A). The number of metastases and tumor sizes varied among individuals, and in some extreme cases large tumors covered most of the lungs. The NTF2 high dox+ mice generally developed fewer and smaller tumors that covered less lung area (Fig. 3B). Tissue staining with Melan A and S100 confirmed that the tumors were human melanoma-derived. Although metastases were primarily found on the lungs, some NTF2 low mice also exhibited tumors on the spleen and spinal cord, manifesting with paralyzed hind legs and enlarged bladders. We also quantified statistically significant differences in lung weights between NTF2 low and NTF2 high dox+ (Fig. 3C).

**Figure 3:**
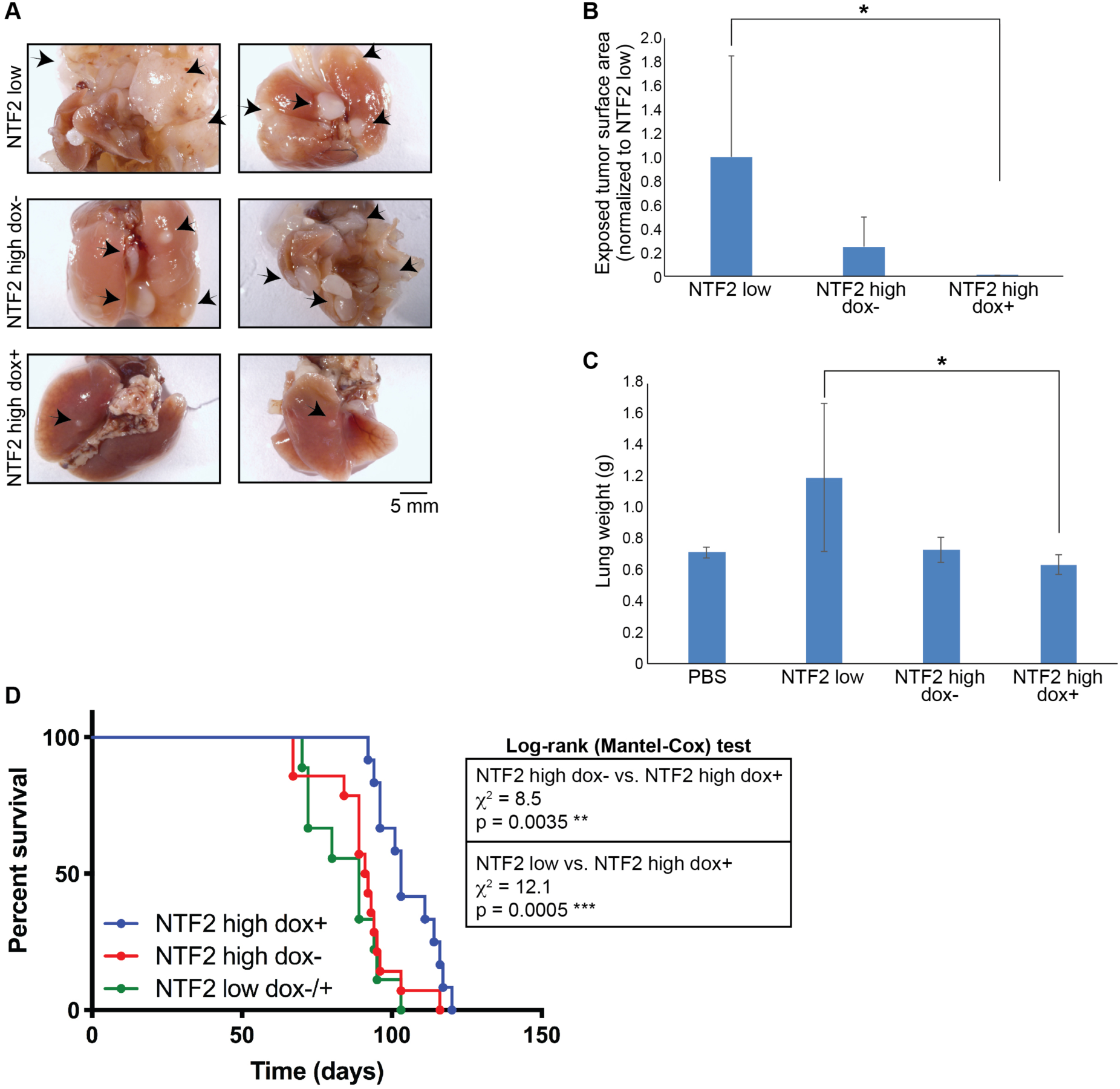
Increasing NTF2 expression in metastatic melanoma reduces lung metastasis and prolongs survival. **(A-C)** Rag2^−/−^ γc^−/−^ knockout mice (B6) were injected with NTF2 low or NTF2 high melanoma cells and fed either normal food or food containing 200 mg/kg doxycycline. Mice injected with NTF2 high melanoma cells and treated with doxycycline are labeled as NTF2 high dox+. **(A)** Representative whole lung images showing metastatic melanoma nodules (black arrows). **(B)** The total area of surface exposed tumors on each lung was measured, averaged, and normalized to NTF2 low. Each group included 3-5 mice, and the experiment was repeated twice. Error bars are SD. * p < 0.05. **(C)** Individual lungs were weighed. The control group includes lungs from mice not injected with tumor cells and mice injected with PBS. Each group included 4-5 mice, and the experiment was repeated twice. Error bars are SD. * p < 0.05. **(D)** Rag2^−/−^ γc^−/−^ knockout mice (BALB/c) were assayed for survival after cell injection. Survival curves are shown for NTF2 low dox -/+ (n=9), NTF2 high dox-(n=14), and NTF2 high dox+ (n=12). The NTF2 low curve includes 3 mice treated with doxycycline and 6 mice not treated with doxycycline. Data were pooled from two independent experiments. Log-rank (Mantel-Cox) tests were performed in GraphPad Prism.

We performed separate survival experiments with Rag2^−/−^ γc^−/−^ knockout mice intravenously injected with NTF2 high cells and fed doxycycline-containing food or normal food. The NTF2 high dox+ mice showed better survival than the NTF2 high dox-mice, living 12 days longer based on median survival times (Fig. 3D). In parallel, we observed that mice injected with NTF2 low cells (dox-/+) died two weeks earlier than NTF2 high dox+ mice. Together these data show that higher NTF2 levels can reduce melanoma metastasis and prolong overall survival of animals with metastatic melanoma.

## Identification of differentially expressed genes in cells expressing different levels of NTF2

We next addressed how increasing NTF2 levels affects gene expression by performing triplicate whole transcriptome sequencing of the NTF2 low and NTF2 high dox+ cell lines. Transcriptomes were compared to identify differentially expressed genes (DEG) between these two cell groups. For the DEG analysis, Wald tests were performed to generate fold-changes in gene expression levels and associated p-values. Our cutoff for DEGs was absolute fold changes > 2 with a false discovery rate (FDR) < 0.05. While one of the NTF2 low samples showed an expression profile quite different from the other two samples, the replicates generally correlated within a given cell group (Fig. 4A). This was further confirmed by two-dimensional principal component analysis (PCA) where the NTF2 low and NTF2 high dox+ samples fell into two distinct groups (Fig. 4B). Among replicates, the NTF2 high dox+ samples showed less variability than the NTF2 low samples.

**Figure 4:**
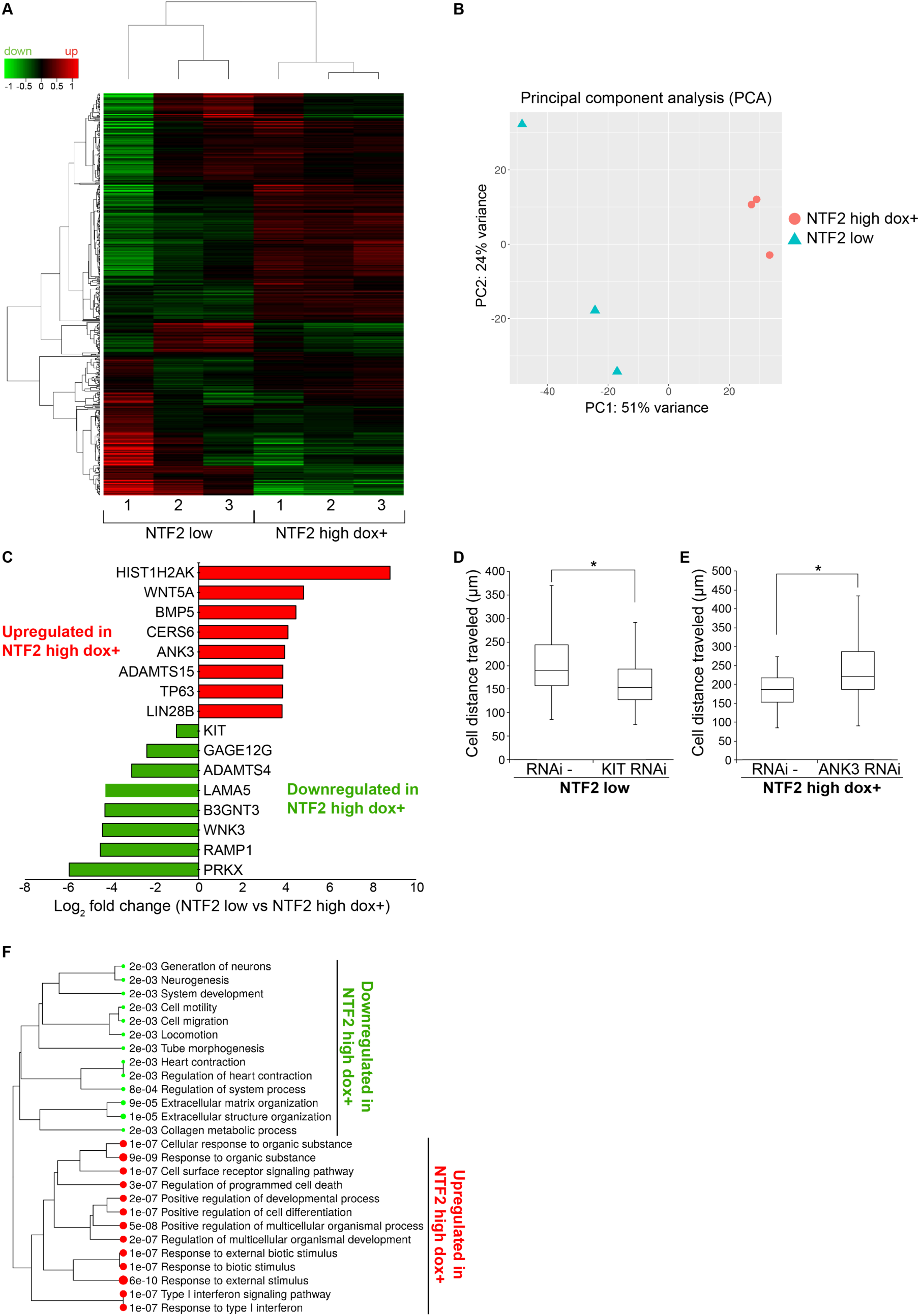
Increasing NTF2 levels alters expression of cancer- and cell migration-related genes. RNAseq data were acquired for three independent samples each of NTF2 low and NTF2 high dox+ cells. Also see Table S1. **(A)** Expression heat map of the 500 most differentially expressed genes with hierarchical clustering. Red and green represent genes upregulated and downregulated, respectively, relative to a reference transcriptome. The Z score cut-off was 4. **(B)** Two dimensional principal component analysis (PCA) is shown. **(C)** Based on log_2_ fold change ranking, the most highly upregulated and downregulated cancer-related genes in NTF2 high dox+ cells are shown. Cutoff: FDR < 0.05, minimum fold change = 2. **(D)** KIT gene expression was knocked down by siRNA in NTF2 low cells, and individual cell migration over 10 hours was assayed and quantified as in Fig. 2C-D for at least 38 cells per condition from two independent experiments. **(E)** ANK3 gene expression was knocked down by siRNA in NTF2 high dox+ cells, and individual cell migration over 10 hours was assayed and quantified as in Fig. 2C-D for at least 30 cells per condition from two independent experiments. **(F)** Enriched pathways are listed for differentially expressed genes. Error bars are SD. * p < 0.05.

We identified 479 genes that were differentially expressed between the NTF2 low and NTF2 high dox+ cell lines. Increased NTF2 expression led to the downregulation of 322 genes and upregulation of 157 genes (Table S1). Among those genes downregulated in NTF2 high dox+, many are associated with tumor promoting characteristics, including cell proliferation (RAMP1, B3GNT3), cell migration (LAMA5, PRKX, KIT), and angiogenesis (PRKX) (Li et al., 2011; Logan et al., 2013; Zhang et al., 2015) (Fig. 4C). The KIT gene plays an important role in melanocyte development and migration and its expression is elevated in advanced melanomas, but downregulated in the NTF2 high dox+ cells. Genes upregulated in NTF2 high dox+ are involved in suppressing tumor growth and invasion (ADAMTS15, ANK3, CERS6) and prolonging patient survival (TP63, BMP5) (Monti et al., 2017; Porter et al., 2006; Romagnoli et al., 2012). For example, while low levels of CERS6 in melanoma are associated with increased cell proliferation and invasion (Tang et al., 2016), CERS6 is upregulated in NTF2 high dox+. In another example, high levels of ANK3 suppress tumor cell invasion (Wang et al., 2016). To test the functional significance of a subset of these DEGs, we performed gene knockdowns. Reducing KIT expression in NTF2 low cells decreased cell migration, while ANK3 knockdown in NTF2 high dox+ cells increased cell migration (Fig. 4D-E), consistent with their previously described roles in cancer.

To obtain a global view of the biological processes and gene families impacted by NTF2 expression levels, we performed gene ontology (GO) enrichment analysis for DEGs (Fig. 4F). Genes upregulated in NTF2 high dox+ are involved in the regulation of programmed cell death (35 genes), positive regulation of cell differentiation (27 genes), and cell surface receptor signaling pathways (51 genes). GO analysis of genes downregulated in NTF2 high dox+ are enriched for cell motility (43 genes), cell migration (42 genes), and extracellular matrix organization (19 genes). GO cellular components analysis showed that genes upregulated in NTF2 high dox+ produce primarily nuclear proteins (e.g. components of nucleosomes, Cajal bodies, chromosomes, and chromatin), while downregulated gene-products are predominantly related to extracellular functions (e.g. components of the extracellular matrix, exocytic vesicles, and plasma membrane).

We also compared the NTF2 low transcriptome to that of NTF2 high dox-. While the PCA analysis showed two distinct groups, we noted that the replicates were more tightly clustered with higher NTF2 expression (Fig. S2). Comparing this PCA to the PCA for NTF2 low versus NTF2 high dox+ (Fig. 4B), we can conclude that replicate variability is reduced as NTF2 expression levels increase. These data are consistent with the idea that NTF2 expression can partially suppress the highly variable gene expression patterns characteristic of cancer cells (Dagogo-Jack and Shaw, 2018). Given their “leaky” NTF2 expression, NTF2 high dox-cells show a gene expression pattern more similar to that of NTF2 high dox+ than to the NTF2 low parental cell line. We constructed Venn diagrams to identify similarities in gene expression profiles among these cell lines. About two-thirds of the same genes are differentially expressed when NTF2 high dox- and NTF2 high dox+ are compared to NTF2 low. We also directly compared the transcriptomes of the NTF2 high dox- and NTF2 high dox+ cell lines (Table S2), using a p-value cutoff < 0.05 because few genes passed the FDR < 0.05 test. 63 genes were upregulated and 260 genes were downregulated in NTF2 high dox+ compared to NTF2 high dox-. Many of these genes correspond to long non-coding RNAs, some having been implicated in cell proliferation and oncogenesis (e.g. MIR155HG and SOX2-OT) (Due et al., 2016; Wang et al., 2017). Many of the differentially expressed protein-coding genes play roles in carcinogenesis (e.g. GAGE12J, VWA2, NUAK1, and MGAT4C), consistent with the transcriptional differences we identified between the NTF2 low and NTF2 high dox+ cell lines (Chen et al., 2013; Demichelis et al., 2012; Gonzalez et al., 2018; Huang et al., 2014).

### NTF2 levels affect lamin A nuclear localization, nuclear size, heterochromatin distribution, and gene positioning

Because NTF2 associates with the NPC and can limit nuclear import of large cargo molecules like nuclear lamins, we next examined if lamin A incorporation into the nuclear lamina differs as a function of NTF2 level in our melanoma cell lines. Based on immunofluorescence, lamin A protein levels at the nuclear envelope were higher in NTF2 low cells compared to NTF2 high dox+ cells (Fig. 5A), consistent with reduced NTF2 at the NPC permitting increased lamin import. Furthermore, nuclear size and lamin A nuclear localization were increased in NTF2 low cells relative to cells with higher NTF2 expression (Fig. 5A-B), consistent with previous reports in *Xenopus* and mammalian cells (Jevtic et al., 2015; Levy and Heald, 2010; Vukovic et al., 2016). We also found that ectopic NTF2 expression reduced nuclear size in LNCaP prostate cancer cells and altered colony morphology (Fig. S1D-E).

**Figure 5:**
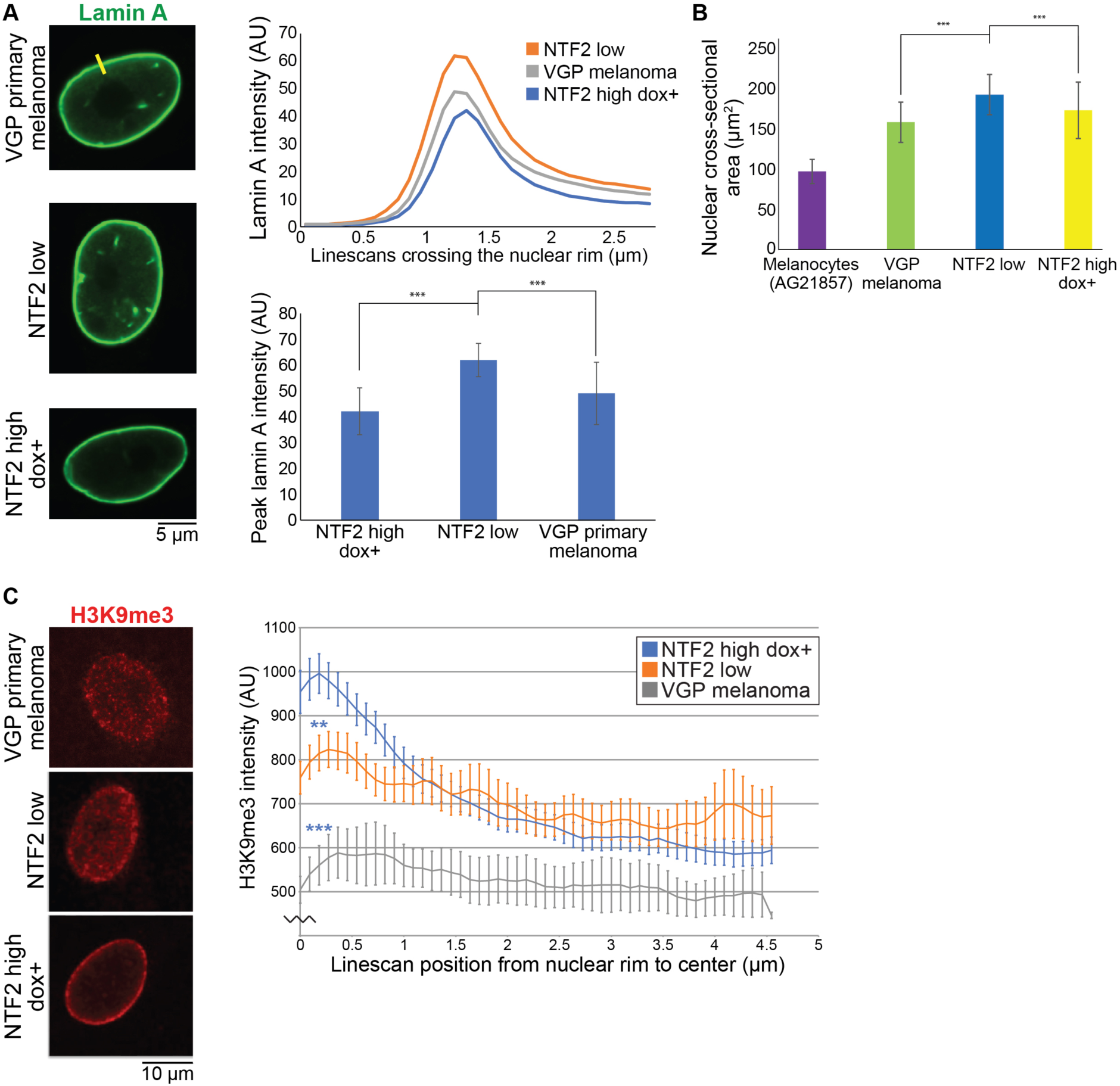
NTF2 affects lamin A levels at the nuclear envelope, nuclear size, and heterochromatin distribution. VGP primary melanoma refers to a primary vertical growth phase melanoma cell line (WM983A) that was derived from the same patient as NTF2 low (WM983B) and represents an earlier stage of disease. **(A)** The indicated cells were stained for lamin A and representative images are shown. Line scans were acquired across the nuclear rim as indicated by the yellow line. In the top graph, lamin A intensity was quantified across at least 50 line scans per cell line for two independent experiments (see Methods). In the bottom graph, the highest lamin A intensities from each line scan were averaged. Error bars are SD. *** p < 0.0001. **(B)** Nuclear cross-sectional area, which serves as a good proxy for nuclear surface area and volume (Edens and Levy, 2014; Jevtic and Levy, 2015; Levy and Heald, 2010; Vukovic et al., 2016), was measured for at least 100 Hoechst-stained nuclei per cell line. The normal melanocyte cell line is AG21857. Error bars are SD. *** p < 0.001. **(C)** The indicated cells were stained for the heterochromatin marker H3K9me3 and representative images are shown. To quantify heterochromatin distribution, H3K9me3 intensity was quantified along line scans drawn from the nuclear rim toward the center (see Methods). Averages from at least 150 line scans are shown for each cell line for two independent experiments. Error bars are SEM. ** p < 0.01, *** p < 0.005. The statistical comparisons are for NTF2 low and VGP primary melanoma compared to NTF2 high dox+ at line scan position 0.2 µm.

Adjacent to the inner nuclear membrane, the nuclear lamina contacts peripheral chromatin through lamina-associated domains that generally repress gene expression, and lamin levels and localization may regulate chromatin organization and gene expression (Lochs et al., 2019; van Steensel and Belmont, 2017). Furthermore, Gene Set Enrichment Analysis identified chromatin/gene silencing and nucleosome assembly pathways as being upregulated in NTF2 high dox+ cells, consistent with the large number of downregulated genes in these cells. For these reasons, we examined the heterochromatin distribution in our cell lines. While H3K9me3 heterochromatin was fairly evenly distributed throughout the nucleus in NTF2 low cells, we observed a striking increase in the H3K9me3 signal at the nuclear envelope in NTF2 high dox+ cells (Fig. 5C).

Given that NTF2 levels affect heterochromatin distribution and gene expression, we wanted to determine if the intranuclear positioning of DEGs might differ in our cell lines. To visualize gene positions we performed DNA fluorescent *in situ* hybridization (FISH) on the following genes: KIT, PADI2, and RAMP1. KIT is often overexpressed in melanoma, where it activates several signaling pathways and increases cell proliferation and metastasis (Curtin et al., 2006; Willmore-Payne et al., 2006). Our transcriptomics data showed that KIT gene expression is low in VGP primary melanoma and NTF2 high dox+ cells but highly expressed in NTF2 low cells. Based on FISH, we found that the KIT gene is generally close to the nuclear periphery in NTF2 low cells, as opposed to its more central nuclear location in NTF2 high dox+ cells and VGP primary melanoma cells (Fig. 6A). In NTF2 low cells, the PADI2 gene is centrally located within the nucleus and its expression level is high, while in NTF2 high dox+ cells PADI2 re-localizes to the nuclear envelope and is downregulated (Fig. 6B). To determine if lamin A levels affect gene positioning in our cells, we transiently overexpressed lamin A in NTF2 high dox+ cells and performed FISH for the RAMP1 gene. We found that ectopic lamin A expression caused RAMP1 to re-localize from the nuclear periphery more toward the center of the nucleus (Fig. 6C). Taken together, we observed a correlation between the expression level and nuclear positioning of several genes, suggesting that NTF2 might impact gene expression by influencing gene positioning, potentially through altered lamin A nuclear localization and/or nuclear size.

**Figure 6:**
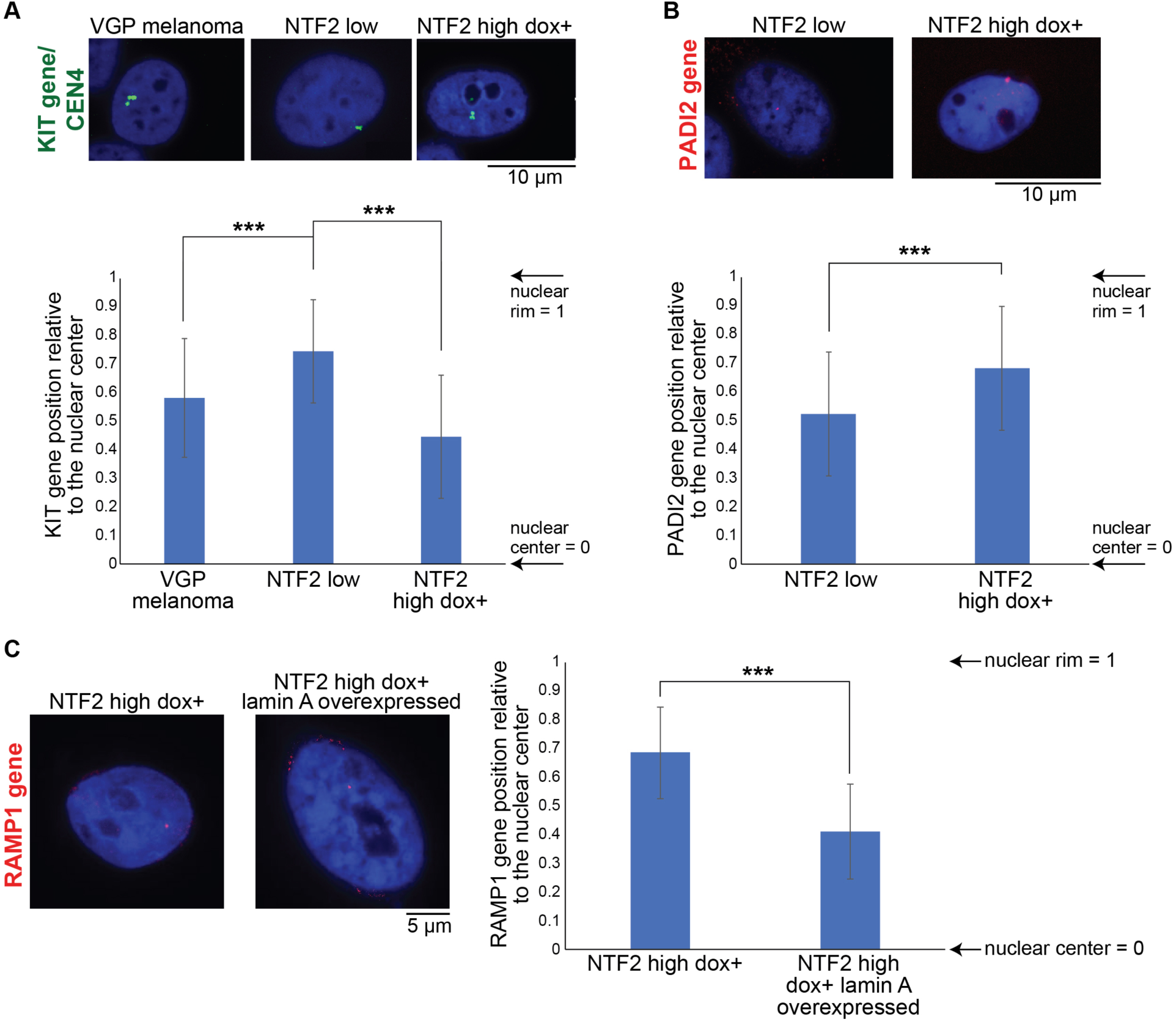
NTF2 levels affect gene positioning. **(A)** DNA FISH was performed for the KIT gene and the centromere of chromosome 4 (CEN4). The two signals overlapped and representative CEN4 images are shown due their stronger signal. Gene distance from the nuclear center was divided by the nucleus radius (see Methods). At least 30 nuclei were measured for each condition. **(B)** DNA FISH was performed for the PADI2 gene and representative images are shown. Gene distance from the nuclear center was divided by the nucleus radius (see Methods). At least 30 nuclei were measured for each condition. **(C)** NTF2 high cells were transiently transfected to overexpress lamin A. Control and transfected cells were treated with 20 ng/µl doxycycline for 48 hours. DNA FISH was then performed for the RAMP1 gene and representative images are shown. Gene distance from the nuclear center was divided by the nucleus radius (see Methods). At least 30 nuclei were measured for each condition. Error bars are SD. *** p < 0.0001.

### Increasing NTF2 expression in metastatic melanoma produces cells with primary melanoma-like characteristics

Since increasing NTF2 expression in metastatic melanoma reduced nuclear size, altered gene positioning and expression, decreased cell motility and proliferation, and increased apoptosis, we conclude that certain cancer cell phenotypes can be abrogated by ectopic NTF2 expression. For this reason, we hypothesized that NTF2 high dox+ cells might be more similar to VGP primary melanoma cells (WM983A) than to the original NTF2 low metastatic melanoma cells (WM983B) from which they were derived. NTF2 levels are 1.9 ± 0.8 (average ± SD) fold higher in the VGP primary melanoma cells compared to NTF2 low metastatic melanoma cells, consistent with our previously published work showing an inverse correlation between NTF2 expression levels and cancer progression (Vukovic et al., 2016). As predicted by these differences in NTF2 expression levels, VGP primary melanoma nuclei are smaller than nuclei in NTF2 low cells and similar in size to NTF2 high dox+ nuclei (Fig. 5B).

We compared the transcriptomes of triplicate samples of VGP primary melanoma, NTF2 low, and NTF2 high dox+. Heat maps, PCA analyses, and volcano plots showed clustering of the cell lines into three separate groups, with the VGP primary melanoma being the most distinct (Fig. 7A-C). 43% of the genes upregulated in NTF2 high dox+ cells were also upregulated in VGP primary melanoma when compared to the metastatic parental cell line NTF2 low (Table S3). Similar pathways associated with DEGs were identified when comparing NTF2 low to both VGP primary melanoma and NTF2 high dox+ (Fig. 7D and 4F). Furthermore, Reactome pathway analysis implicated extracellular matrix degradation, collagen degradation, and laminin interaction pathways more strongly in NTF2 low cells than in VGP primary melanoma or NTF2 high dox+ (Fig. 7E). These data suggest that a similar subset of genes is impacted by ectopic NTF2 expression in metastatic melanoma and in the progression from VGP primary melanoma to metastatic melanoma. Consistent with this idea, we noted that KIT gene localization was similar in NTF2 high dox+ cells and VGP primary melanoma cells (Fig. 6A). Lastly, cell migration assays showed that VGP primary melanoma cells migrate more slowly than NTF2 low cells and more similarly to NTF2 high dox+ cells (Fig. 7F and 2D). Taken together, these data suggest that increasing NTF2 expression levels in metastatic melanoma can lead to at least partial reversion to an earlier stage of disease.

**Figure 7:**
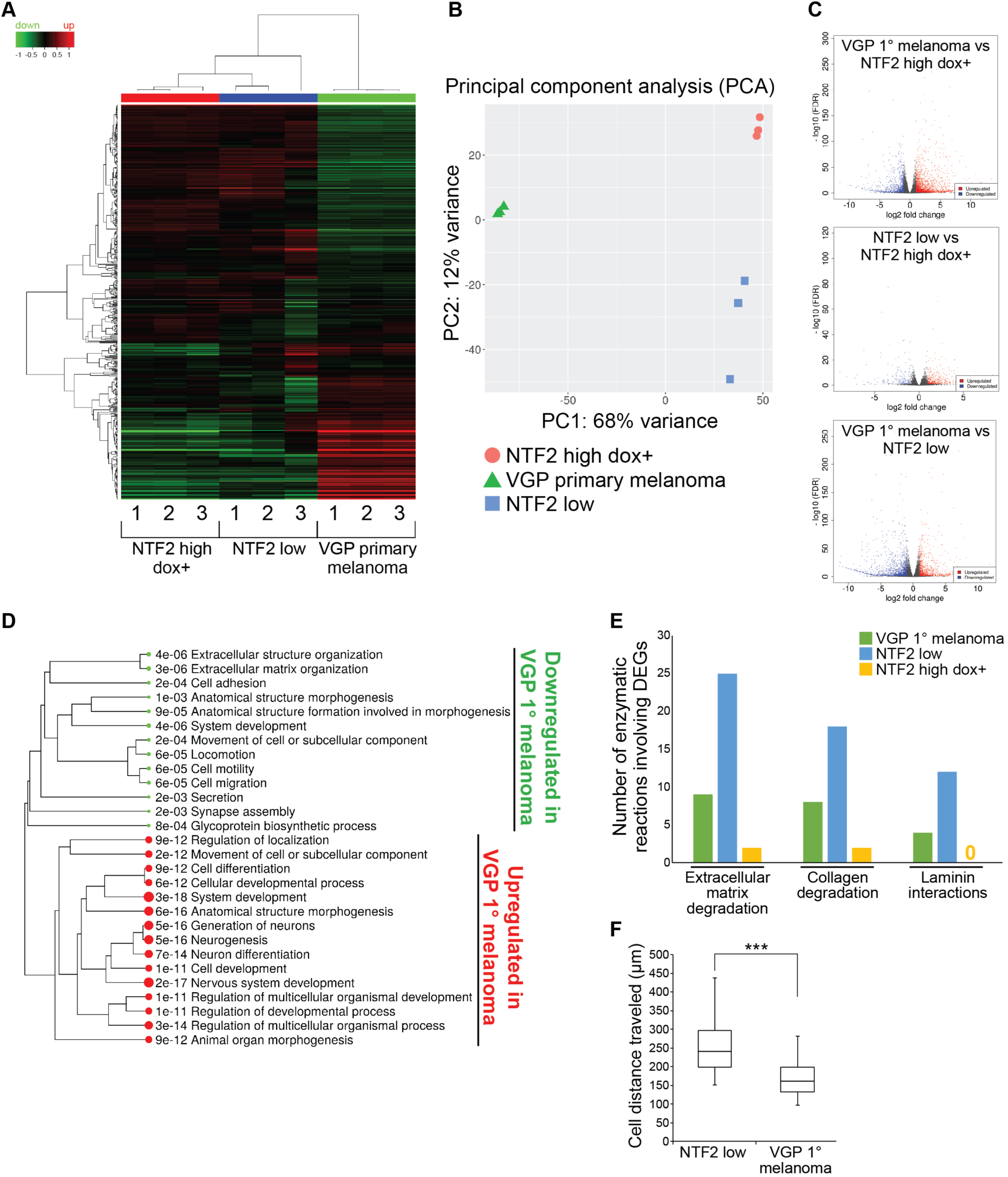
Increasing NTF2 expression in metastatic melanoma generates a gene expression pattern with characteristics of primary melanoma. RNAseq data were acquired for three independent samples each of NTF2 low, NTF2 high dox+, and VGP primary melanoma cells. The NTF2 low and NTF2 high dox+ data are the same shown in Fig. 4. Also see Table S3. **(A)** Expression heat map of the 1000 most differentially expressed genes with hierarchical clustering. Red and green represent genes upregulated and downregulated, respectively, relative to a reference transcriptome. The Z score cut-off was 4. **(B)** Two dimensional principal component analysis (PCA) is shown. **(C)** Volcano plots show various pairwise comparisons in gene expression. Upregulated and downregulated genes are red and blue, respectively. **(D)** Enriched pathways are listed for genes that are differentially expressed between VGP primary melanoma and NTF2 low cells. **(E)** Results of Reactome pathway analysis are shown (see Methods). **(F)** Individual cell migration assays were performed and quantified as in Fig. 2C-D. At least 30 cells were tracked per cell line. Error bars are SD. *** p = 0.0002. Note that the “NTF2 low” data shown here are the same as in Fig. 2D.

## DISCUSSION

In this study we present new cellular roles for NTF2 that are relevant to cancer progression. While NTF2 was previously known to function in nucleocytoplasmic transport, Ran recycling, and nuclear size regulation, we now show that NTF2 levels influence a number of cancer cell phenotypes in melanoma. During melanoma progression, there is a general reduction in NTF2 levels (Vukovic et al., 2016). Increasing NTF2 expression in metastatic melanoma slowed cell motility to that observed in primary melanoma, reduced metastatic potential and cell proliferation, and increased apoptosis. Furthermore, our transcriptomics analysis revealed similarities in the gene expression profiles of metastatic melanoma cells expressing higher NTF2 levels and VGP primary melanoma cells. These data suggest that increasing NTF2 expression is sufficient to at least partially revert metastatic melanoma cells to a more primary melanoma character, supporting the idea that NTF2 is a melanoma tumor suppressor and potential new therapeutic target to treat metastatic melanoma. Open questions remain about the mechanisms that account for reductions in NTF2 levels during progression from normal melanocytes to VGP primary melanoma to metastatic melanoma. There may be changes in the transcriptional program that regulates NTF2 expression and/or acquisition of NTF2-destabilizing mutations. Another possibility is that the tumor microenvironment plays a role in epigenetically altering NTF2 expression during cancer progression (Barcellos-Hoff, 2001; Molognoni et al., 2011; Seftor et al., 2005; Villanueva and Herlyn, 2008; Wu et al., 2017). Our study also raises questions about additional factors that regulate nuclear architecture and their role in, not only melanoma, but also other cancers.

How might NTF2 levels influence the various cancer cell characteristics examined in this study? It is now well established that chromosomes occupy specific positions within the nucleus, so-called chromosome territories, that in turn influence gene expression (Cremer and Cremer, 2010; Finn and Misteli, 2019). One possibility is that NTF2-induced changes in nuclear size cause chromosomes and genes to re-position. It is also known that specific chromosomal regions interact with the nuclear lamina, forming lamina-associated domains (LADs) that generally correspond to heterochromatic silenced regions of the genome (van Steensel and Belmont, 2017). Because the amount of lamin A at the NE is sensitive to NTF2 levels, another model is that LADs are altered when NTF2 expression is manipulated. In either case, the prediction is that NTF2 levels influence gene positioning and expression, both results borne out in our DNA FISH, heterochromatin localization, and transcriptomics data. Of note, increasing NTF2 expression upregulated pathways involved in gene silencing, nucleosome assembly and organization, and negative epigenetic gene regulation. Thus, we propose that increasing NTF2 expression in metastatic melanoma reduces lamin A import and nuclear size, leading to altered gene positioning and expression that reduce cell proliferation, migration, and metastasis.

Interestingly, another nuclear transport and sizing factor, importin α, has already been suggested for use as a cancer biomarker. Higher importin α expression correlates with poor survival rates in breast cancer (Gluz et al., 2008), and importin α knockdown in non-small cell lung carcinoma suppressed cell proliferation and migration (Wang et al., 2012; Wang et al., 2011). Similarly, it has been shown that changes in importin α levels have prognostic value in many different cancers, including lung, prostate, brain, and liver cancer (Christiansen and Dyrskjot, 2013; Gao et al., 2018; Gousias et al., 2012; Wang et al., 2011). Consistent with these observations, we found that ectopic importin α overexpression in NTF2 high dox+ cells resulted in larger nuclei and higher rates of cell proliferation (data not shown).

Increasing NTF2 levels in metastatic melanoma upregulated expression of a number of genes known to be associated with better survival rates in patients. ANK3, a negative regulator of cell invasion, was one of the most highly expressed genes in NTF2 high dox+ cells and VGP melanoma. ANK3, TP63, and ADAMTS15 were each highly upregulated in NTF2 high dox+ cells and are associated with improved patient survival in certain cancers (Ma et al., 2015; Porter et al., 2006; Wang et al., 2016; Wei et al., 2010). Another highly expressed gene in NTF2 high dox+ cells was BMP5, a gene whose downregulation in breast cancer is associated with disease recurrence (Romagnoli et al., 2012). Thus NTF2 overexpression generates a gene profile associated with improved prognosis in many cancers, consistent with our mouse studies showing that increasing NTF2 levels in metastatic melanoma cells reduces lung metastases and increases survival.

Might increasing NTF2 expression represent a novel approach to cancer treatment? In cultured prostate cancer cells, increasing NTF2 expression reduced cell proliferation and nuclear size while increasing sensitivity to Taxol treatment (Fig. S1 and data not shown). With regard to melanoma, most currently available drugs inhibit BRAF and MAPK signaling (Broman et al., 2019; Gibney and Zager, 2013). Despite some of these drugs being temporarily effective against melanoma, most patients develop drug resistance over time. For example, roughly half of melanoma patients treated with Dabrafenib or Trametinib develop resistance within 6-7 months after starting therapy (Flaherty et al., 2012). One factor known to contribute to resistance is increased expression of the growth factor binding protein IGFBP2 (Strub et al., 2018). Highly expressed in our NTF2 low metastatic melanoma cells, IGFBP2 was downregulated upon NTF2 overexpression, suggesting these cells might show an improved drug response. Given that combination therapy often leads to better outcomes, we propose the exploration of approaches to increase NTF2 expression in cancer, approaches that may improve patient survival especially when combined with currently available drug treatments.

## MATERIALS AND METHODS

### Cell lines

The human melanocyte cell line (AG21857) was obtained from the NIA Cell Culture Repository through the Coriell Institute. Melanoma cell lines (WM983A and WM983B) were obtained from the Wistar repository through the Coriell Institute. The human prostate cancer cell line (LNCaP) was obtained from Jay Gatlin (University of Wyoming). AG21875 cells were maintained in Medium 254 (#M254500, Invitrogen) supplemented with HMGS (#S0025, Invitrogen). WM983A and WM983B cells were maintained in 4:1 MCDB153 (#M7403, Sigma-Aldrich) and Leibovitz L-15 (#L4386, Sigma-Aldrich), supplemented with 5 µg/ml insulin (#I9278, Sigma-Aldrich), 2% FBS, 1.68 mM CaCl_2_, and 50 IU/ml penicillin and streptomycin. LNCaP cells were maintained in RPMI-1640 (Sigma) supplemented with 7.5% FBS, and 50 IU/ml penicillin and streptomycin. All cells were grown at 37°C and 5% CO_2_.

### Plasmids

Plasmid pDL23 was described previously and consists of human NTF2 cloned into pEmCherry-C2 (a derivative of pEGFP-C2, a gift from Anne Schlaitz) (Vukovic et al., 2016). The mCherry-NTF2 fragment was excised from pDL23 at AgeI/BclII and cloned into pTetOne (#634303 Clontech, Takara) at AgeI/BamHI to generate pTetOne mCherry-NTF2 (pDL65). Plasmid pDL65 was digested with EcoRI to remove mCherry and recircularized to generate pTetOne NTF2 (pDL66). Plasmid pTet-tTs was from Clontech (#631011). A plasmid consisting of human lamin A cloned into pcDNA3.1(+) was described previously (pDL80) (Jevtic et al., 2015). Human importin α (hSRP1α) was cloned from pKW230 (a gift from Karsten Weis) into pEmCherry-C2 at EcoRI/KpnI to generate pEmCherry-C2 importin α (pDL21).

### NTF2-inducible melanoma cells

WM983B cells were seeded into 6-well plates and transfected at 70% confluency using Xfect Transfection Reagent (#631317, Clontech) with 5 µg of pTetOne NTF2 (pDL66) and 100 ng of linear hygromycin marker (#631625, Clontech). After 4 hours at 37°C, the media was replaced. 48 hours post transfection, confluent cells were split into 4 x 10 cm dishes. 96 hours post transfection, selective antibiotic hygromycin (#631309, Clontech) was added at 200 µg/ml. Antibiotic-resistant colonies appeared after two weeks. Each colony was transferred to a well of a 24-well plate to expand and test for NTF2 expression. To screen for NTF2 induction, clonal cell populations were treated with 1, 5, 10, 20, 50, 100, or 1000 ng/ml of doxycycline (#631311, Clontech), and NTF2 levels were assayed by western blot after 24 and 48 hours.

### NTF2-overexpressing prostate cancer cells

LNCaP cells were seeded into 6-well plates and transfected at 70% confluency using Xfect Transfection Reagent (#631317, Clontech) with 5 µg of pEmCherry-C2 NTF2 (pDL23). After 4 hours at 37°C, the media was replaced. 48 hours post transfection, confluent cells were split into 4 x 10 cm dishes. 96 hours post transfection, selective antibiotic geneticin was added at 500 µg/ml. Antibiotic-resistant colonies appeared after two weeks. Each colony was transferred to a well of a 24-well plate to expand and test for NTF2 expression.

### Transient transfections

Cells were grown to 70-80% confluency and transient transfections were performed using Lipofectamine 3000 (Invitrogen, L3000008). In one tube, 2.5 µg of DNA were added to 125 µL of OptiMEM media plus 5 µL of P3000 reagent. In another tube, 5 µl of Lipofectamine 3000 reagent was added to 125 µL of Opti-MEM media. The contents of both tubes were mixed and incubated for 5 minutes at room temperature. 250 µL of DNA-lipofectamine complex were added to each well of a 6-well plate. The media was changed after 24h, and cells were fixed and visualized 48h post-transfection. Transient RNAi transfections were performed using Lipofectamine RNAiMAX (Invitrogen, 13778075). Briefly, 3 µl of 10 µM siRNA were diluted in 50 µl of Opti-MEM media, 3 µl of Lipofectamine RNAiMAX reagent were diluted in 50 µl of Opti-MEM media, and the two were mixed and incubated for 5 minutes at room temperature. The 100 µl transfection mixture was added to 500 µl of culture media and incubated for 48h. The following RNAi constructs were used: ANK3 (#187683817, IDT), KIT (#187683808, IDT), transfection control TYE 563-labeled DsiRNA (#51-01-20-19, IDT). After transient transfections, cells on coverslips were washed with PBS (without Mg^2+^ and Ca^2+^), fixed with 4% paraformaldehyde for 20 minutes, washed, stained with 5 µg/ml Hoechst for 5 minutes, washed, mounted in Vectashield (Vector Laboratories), and sealed with nail polish.

### Immunofluorescence

Cells grown on glass coverslips or in chamber slides (Thermo Fisher Scientific, #177437PK) were washed twice with PBS, fixed with 4% paraformaldehyde for 20 min at room temperature, and then subjected to three 5-min washes with PBS. Cells were permeabilized for 10 min with 0.5% Triton X-100 in PBS and washed three times with PBS for 5 min each. Blocking was performed for 40 min at room temperature with 5% BSA and 0.05% Tween 20 in PBS, and then incubated overnight at 4°C with primary antibody diluted in 1% BSA in PBS. Cells were washed three times in PBS plus 0.05% Tween 20 for 5 min each and incubated at room temperature for 1 h with secondary antibodies diluted in 1% BSA in PBS. Cells were then stained with 5 µg/ml Hoechst in PBS for 5 min and washed three times in PBS plus 0.05% Tween 20 for 5 min each. Coverslips were mounted in Vectashield (Vector Labratories) and sealed with nail polish. Primary antibodies included: H3K9me3 monoclonal mouse antibody (Abgent, ADN10307) used at 1:200 and lamin A/C monoclonal mouse antibody (Santa Cruz Biotechnology, sc-376248) used at 1:1000. Secondary antibodies included: Alexa Fluor 488 and Alexa Fluor 568 conjugated goat anti-mouse IgG (Thermo Fisher Scientific, A-11001 and A-11004) used at 1:500.

### Western blots

Whole cell lysates were prepared using SDS-PAGE sample buffer supplemented with benzonase nuclease (Sigma, E1014) and boiled for 5 minutes. Proteins were separated on SDS-PAGE gels (4-20% gradient) and transferred to PVDF membrane. Membranes were blocked in Odyssey PBS Blocking Buffer (Li-Cor, 927-40000). The primary antibodies used were mouse anti-NTF2 at 1:500 (Sigma, N9527), rabbit anti-NTF2 at 1:200 (Aviva System Biology, ARP64840_P050), and mouse anti-actin at 1:200 (Abgent, AM1965b). The secondary antibodies were IRDye 680RD anti-mouse used at 1:20,000 (Li-Cor 925-68070) and IRDye 800CW anti-rabbit used at 1:20,000 (Li-Cor 925-32211). Blots were scanned on a Li-Cor Odyssey CLx instrument and band quantification was performed with ImageStudio. For a given sample, NTF2 band intensity was normalized to the actin signal. For western blots on LNCaP cell lysates, HRP detection was used as previously described (Edens and Levy, 2014).

### Animal study

All mouse (*Mus musculus*) procedures and studies were conducted in compliance with the US Department of Health and Human Services Guide for the Care and Use of Laboratory Animals. Protocols were approved by the University of Wyoming Institutional Animal Care and Use Committee (DHHS/NIH/OLAW Assurance D16-00135, #A-3216-01). Mice were housed in specialized microisolator cages under specific pathogen-free conditions at the University of Wyoming Animal Facility. Mice were maintained in 12 hour light/dark cycles. 6-8 week old female Rag2^−/−^ γc^−/−^ knockout mice in the B6 background (#4111, Taconic Biosciences, Inc.) were used for metastasis assays. 8-10 week old female Rag2^−/−^ γc^−/−^ mice in the BALB/c background (#014593, Jackson Laboratory) were used for survival assays. Mice were fed autoclavable Teklad Global 18% Protein Rodent Diet (#2018S, Envigo). For NTF2 induction, doxycycline hyclate (200 mg per kg) was added to the base diet (En#TD.00502, Envigo). Metastasis and survival assays were performed after intravenous injection of 2.5 x 10^5^ NTF2 low or NTF2 high dox-melanoma cells in 100 µl physiological sterile PBS with 10% OptiMem. For metastasis assays, mice were sacrificed around 4 months after injections to examine metastasis formation. For survival assays, mice were inspected daily until death.

### Mouse tissue collection

Mice were euthanized with isoflurane followed by cervical dislocation. Organs were manually dissected, washed in PBS, fixed overnight in 10% neutral buffered formalin (HT501128, Sigma-Aldrich), and preserved in 70% ethanol. For histology, tissue was embedded in paraffin and 5 mm sections were cut using a microtome. Hematoxylin and eosin tissue staining was performed using HAE-2-IFU (ScyTek Laboratories, Inc). Briefly, slides were deparaffinized in xylene, followed by sequential incubations in 100%, 95%, and 70% ethanol. Tissue sections were stained with Hematoxylin, Mayer’s (Lillie’s Modification) for 5 minutes, washed with two changes of distilled water, incubated with Bluing Reagent for 10-15 seconds, washed with two changes of distilled water, washed with ethanol, stained with Eosin Y Solution (Modified Alcoholic) for 2-3 minutes, and washed with three changes of ethanol. Some of the histology work was performed as a fee-for-service by Dr. Jonathan Fox (Department of Veterinary Sciences, University of Wyoming).

### Cell migration assays

Would healing migration assays were performed. Briefly, 5 x 10^5^ cells were seeded on collagen-coated, chambered coverslips (Ibidi, #80286). After 24h when cells reached confluency, pipette tips were used to generate scratches, 10 µl tips for narrow scratches and 200 µl tips for wider scratches. After changing the media, time lapse confocal imaging was performed for 12-15h, imaging every 5 minutes. Culture conditions were maintained at 37°C and 5% CO_2_ using an INU series Microscope Stage Top Incubator (Tokai Hit; provided by Dr. John Oakey, University of Wyoming). Movies were compiled with NIH ImageJ software (National Institute of Health Bethesda, MD, USA). For narrow scratches, individual cells were tracked using the MTrackJ plugin in ImageJ (Meijering et al., 2012), using the cell nucleus as the central tracking point to measure the total distance traveled. For wide scratches, the scratch area was manually quantified using MetaMorph software.

### Cell proliferation assays

2×10^5^ cells were initially plated in 6-well plates. Every 24h, cells from one plate were trypsinized for 5 minutes, stained with trypan blue, and counted using a Countess Automated Cell Counter (Invitrogen; provided by Dr. Don Jarvis, University of Wyoming).

### Apoptosis assays

For apoptosis assays, cells were incubated for 30 minutes with 40 µl/ml CellEvent™ Caspase-3/7 Green ReadyProbes Reagent (#R37111, Life Technologies). Cells were washed with PBS (without Mg^2+^ and Ca^2+^), fixed with 4% paraformaldehyde for 20 minutes, washed, stained with 5 µg/ml Hoechst for 5 minutes, washed, mounted in Vectashield (Vector Laboratories), and sealed with nail polish. Green apoptotic cells were counted and presented as a percent of the total number of cells. For each coverslip, approximately 500-1400 cells were examined.

### Fluorescence *in situ* hybridization (FISH)

FISH was performed following established protocols provided by Agilent Technologies, with slight modifications. Briefly, cells grown on chambered coverslips were washed three times with PBS, fixed with 4% paraformaldehyde for 20 min, washed three times with PBS for 5 min each, permeabilized with 0.1% Triton X-100 for 5 min, and washed three times with PBS for 5 min each. Cells were then incubated in boiling SSC buffer (15557-044, Invitrogen) for 1 minute in an 1100-watt microwave, followed by a 10-minute incubation in SSC buffer maintained just below boiling in the microwave. Slides were washed with Dako Wash buffer (K5499, Dako) and then washed for two minutes each in increasing concentrations of ethanol (70%, 85%, 100%). After air-drying the slides, probes were added and DNA denaturation was promoted by incubation at 72-74°C for 10 minutes. For hybridization, slides were incubated at 37°C for 12 hours in dark, humid chambers, followed by washing with Dako Stringent Buffer (#K5499) at 63°C for 10 minutes. Slides were then washed for one minute each in increasing concentrations of ethanol (70%, 80%, 100%), dried, and mounted in Dako Florescence Mounting Medium (#K5499) containing 20 µg/ml Hoechst.

The following FISH probes were acquired from Agilent Technologies: SureFISH 4q12KIT 155kb red for the KIT gene (#G101058R, Dako), SureFISH chr4 CEP 613kb green for the centromere of chromosome 4 (#G101066G, Dako), FISH RAMP1 red for the RAMP1 gene (#G110996X-8, Agilent Technologies), SureFISH chr2 CEP 448kb green for the centromere of chromosome 2 (#G101089, Dako), SureFISH PADI2 red for the PADI2 gene (#G110996X-8, Agilent Technologies), SureFISH chr1 CEP 541kb green for the centromere of chromosome 1 (#G101064G, Dako). Prior to use, 1 µl of each probe provided by Agilent was mixed with 8 µl IQFISH Fast Hybridization Buffer (#G9415A, Agilent Technologies, Inc).

### RNA sequencing

RNA extraction, RNA library preparation, and RNA-sequencing were performed as a fee-for-service by GENEWIZ, Inc. For each cell line, frozen cell pellets were provided in triplicate to GENEWIZ, Inc and sequencing was performed using an Illumina HiSeq2500 platform in 2×100bp paired-end configuration in High Output mode (V4 chemistry). Bioinformatics analysis was performed by GENEWIZ, Inc using CLC Genomic Workbench software (Qiagen): sequence quality control, trimming low quality bases and adapter sequences, mapping reads to the genome, determining the read density on genes/exons and annotating, calculating total hit counts and RPKM values for transcripts/genes, and comparing transcript expression levels. Fold changes in expression levels and p-values were generated using the Wald test. For each comparison, genes with a false discovery rate (FDR) < 0.05 and absolute fold-change > 2 were considered as differentially expressed genes. For comparing NTF2 high dox- and NTF2 high dox+, a p-value < 0.05 was used as the cutoff. Heat maps were generated with iDEP.80 (http://bioinformatics.sdstate.edu/idep/) using the heatmap.2 function. Principal component analysis was performed with iDEP.80. Gene ontology enrichment analysis for DEGs was performed with iDEP.80 for biological processes and for cellular components. Pathway analysis was performed with iDEP.80, which takes into account fold-change values for all genes and is independent of DEGs. For pathway analysis, the significance cutoff was a FDR of 0.2, and Gene Set Enrichment Analysis was performed using the preranked mode in the fgsea package. Pathway analysis was also performed using the Reactome Pathway Database (https://reactome.org/) in order to determine the number of genes involved in specific pathways.

### Microscopy and image quantification

For wide-field microscopy, nuclei were visualized using an Olympus BX51 fluorescence microscope with the following Olympus objectives: UPLFLN 40× (NA 0.75, air) and UPLANAPO 60× (NA 1.20, water). Images were acquired with a QIClick Digital charge-coupled device camera, mono, 12-bit (QIClick-F-M12) and cellSens software (Olympus). Cross-sectional nuclear areas were measured from the original thresholded images using MetaMorph software (Molecular Devices). For each coverslip, at least 50, and usually >800, nuclei were quantified for nuclear size measurements. Colony morphology images were acquired with an Olympus SZX16 research fluorescence stereomicroscope, equipped with Olympus DP72 camera, 11.5x zoom microscope body, and SDFPLAPO1XPF objective. Confocal imaging was performed on a spinning-disk confocal microscope based on an Olympus IX71 microscope stand equipped with a five line LMM5 laser launch (Spectral Applied Research) and Yokogawa CSU-X1 spinning-disk head. Confocal images were acquired with an EM-CCD camera (ImagEM, Hamamatsu) using the following Olympus objectives: UPLFLN 10× (NA 0.50, air) and UPLSAPO 60x (NA 0.85, oil). Z-axis focus was controlled using a piezo Pi-Foc (Physik Instrumentes), and multiposition imaging was achieved using a motorized Ludl stage. Generally z-stacks were acquired with a z-slice thickness of 0.2 µm. Image acquisition and all system components were controlled using Metamorph software. Images were acquired with the same exposure time for fluorescence intensity measurements. Heterochromatin H3K9me3 staining distribution was quantified in ImageJ by drawing three straight lines per nucleus (4.5 µm in length) from the nuclear rim to the nuclear center, perpendicular to the nuclear envelope, and measuring pixel intensities along the lines. Pixel intensities along the line scans were then averaged for at least 50 nuclei from two independent experiments. Lamin A staining distribution was quantified in ImageJ by drawing a line (2.7 µm in length) perpendicular to the nuclear envelope, and measuring pixel intensities along the line. Pixel intensities along the line scans were then averaged for at least 50 nuclei from two independent experiments. For FISH experiments, the z-slice with maximum signal intensity was selected. The centroid measurement function in ImageJ was used to determine the center of the nucleus. A line was drawn from the nucleus center to the nucleus rim passing through the FISH signal. The distance between the nucleus center and FISH signal was measured, and the distance between the nucleus center and rim was defined as the nucleus radius. The distance between the gene signal and nucleus center was divided by the nucleus radius, such that a value of 0 or 1 indicates the gene is located at the nucleus center or rim, respectively. For publication, images were cropped and pseudocolored using ImageJ, but were otherwise unaltered.

### Statistics

Averaging and statistical analysis were performed for independently repeated experiments. Two-tailed Student’s t-tests assuming equal variances were performed in Excel (Microsoft) to evaluate statistical significance. Log-rank (Mantel-Cox) tests for survival curves were performed in GraphPad Prism. The p-values, number of independent experiments, and error bars are denoted in the Figure Legends.

## ACKNOWLEDGEMENTS

We thank Dr. Nicolas Blouin (INBRE, University of Wyoming) for bioinformatics assistance, Dr. Jay Gatlin (University of Wyoming) for use of his tissue culture room, Dr. John Oakey (University of Wyoming) for use of his microscope stage top incubator, Dr. Jonathan Fox (University of Wyoming) for mouse tissue analysis, Dr. Wei Guo (University of Wyoming) for use of equipment to prepare tissue slides, and Osimanjiang Wupu (University of Wyoming) for help with mouse spinal cord isolations. The authors declare no competing financial interests. This work was supported by the National Institutes of Health/National Institute of General Medical Sciences (R01GM113028 and P20GM103432) and the American Cancer Society (RSG-15-035-01-DDC).

## AUTHOR CONTRIBUTIONS

Conceptualization, L.D.V., D.L.L.; Investigation, L.D.V. (melanoma studies), K.H.W. (prostate cancer studies), J.P.G. (mouse husbandry and injections); Writing – Original Draft, L.D.V., D.L.L.; Writing – Review & Editing, L.D.V., K.H.W., J.P.G., D.L.L.; Funding Acquisition, D.L.L.; Supervision, D.L.L.

## SUPPLEMENTAL FIGURES

**Figure S1:**
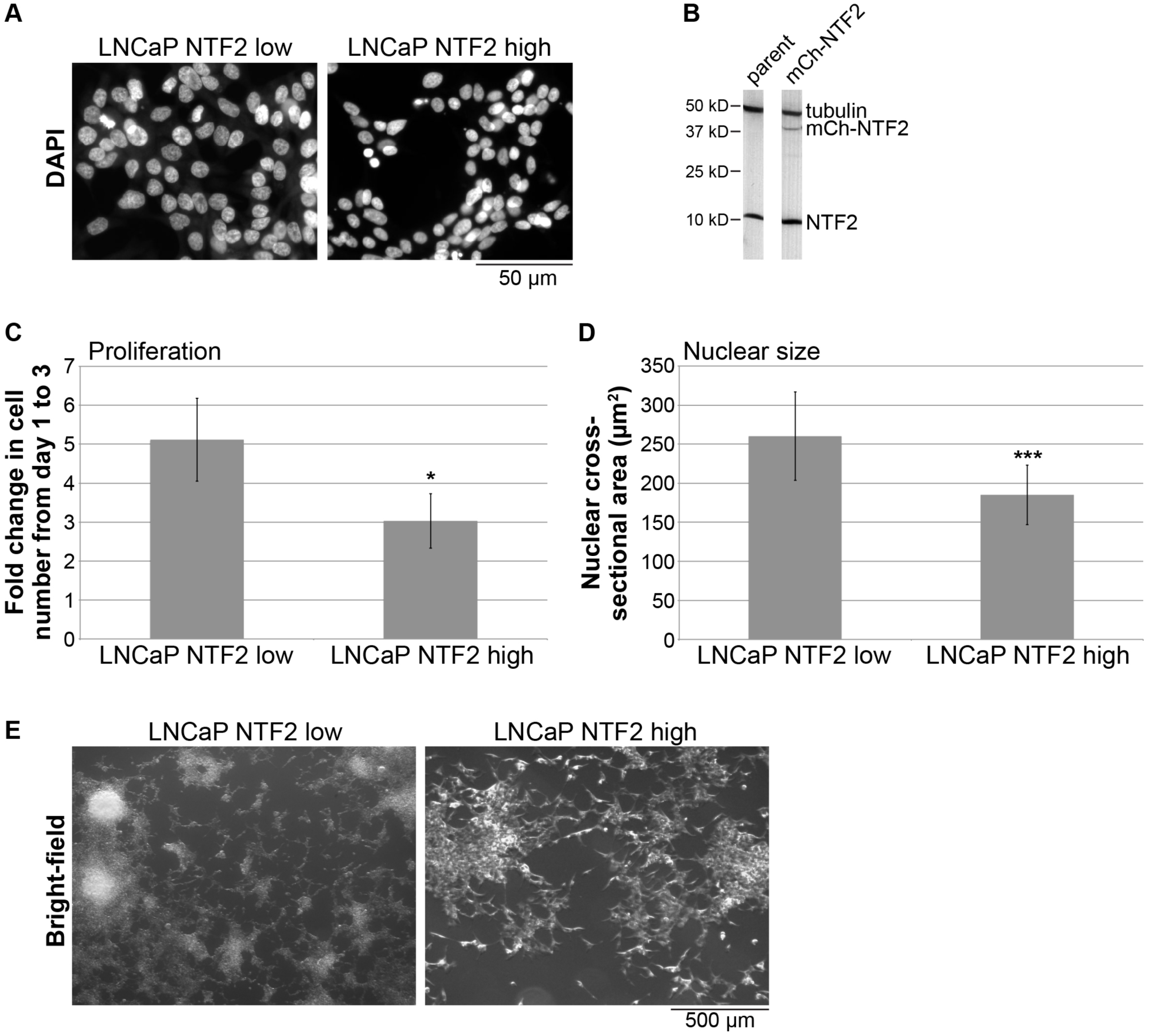
Increasing NTF2 expression in prostate cancer cells decreases cell proliferation and nuclear size. **(A)** LNCaP cells were stably transfected with a construct constitutively expressing mCherry-NTF2. For simplicity, the parent cell line is labeled “LNCaP NTF2 low” and the transfected cell line is labeled “LNCaP NTF2 high.” Representative images of DAPI-stained nuclei are shown. **(B)** Cell lysates from the two cell lines were immunoblotted for NTF2 and tubulin. Ectopic expression of mCherry-NTF2 is apparent in the LNCaP NTF2 high cell line. One representative set of samples is shown out of five. On average, mCherry-NTF2 was overexpressed by 19% ± 7% (average ± SD) over endogenous NTF2 levels. **(C)** Cell proliferation was quantified as described in Fig. 1B. The fold change in cell numbers from day 1 to 3 is plotted for each cell line. **(D)** Cross-sectional nuclear areas were quantified as described in Fig. 5B. More than 150 nuclei were quantified for each cell line. **(E)** Representative bright-field colony morphology images are shown for each cell line. In the parent cell line, cells tend to pile on top of each other in colonies, while this is less apparent in the “LNCaP NTF2 high” cell line. Error bars are SD. * p < 0.05; *** p < 0.001.

**Figure S2:**
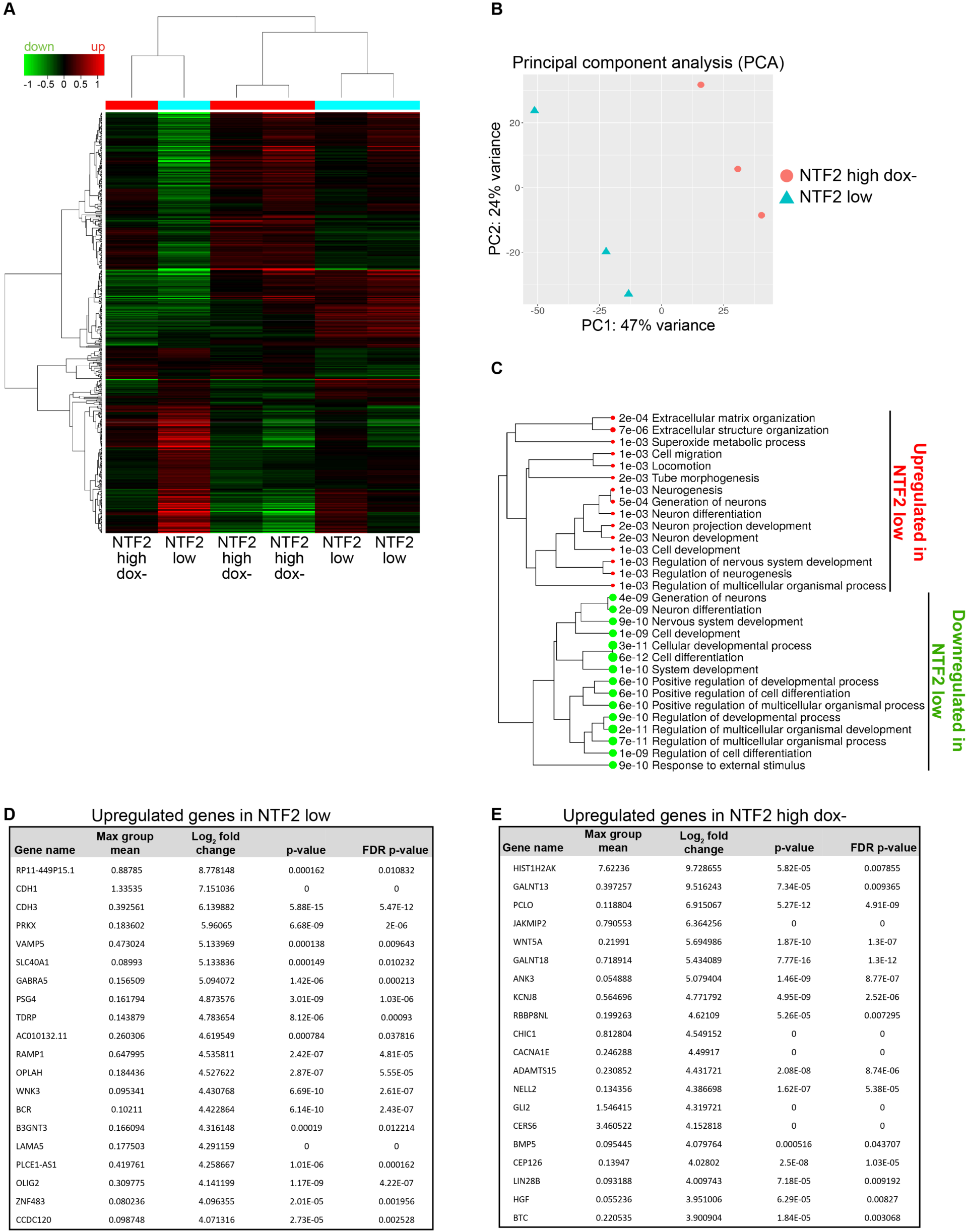
Comparing the transcriptomes of NTF2 low and NTF2 high dox-. RNAseq data were acquired for three independent samples each of NTF2 low and NTF2 high dox-cells. The NTF2 low data are the same described in Fig. 4. **(A)** Expression heat map of the 500 most differentially expressed genes with hierarchical clustering. Red and green represent genes upregulated and downregulated, respectively, relative to a reference transcriptome. The Z score cut-off was 4. **(B)** Two dimensional principal component analysis (PCA) is shown. **(C)** Enriched pathways are listed for differentially expressed genes. **(D)** Based on log_2_ fold change ranking, the most highly upregulated genes in NTF2 low cells are shown. Cutoff: FDR < 0.05. **(E)** Based on log_2_ fold change ranking, the most highly upregulated genes in NTF2 high dox-cells are shown. Cutoff: FDR < 0.05.

## SUPPLEMENTAL TABLE LEGENDS

**Table S1: NTF2 low versus NTF2 high dox+ transcriptomics data.** The RNAseq data are presented for NTF2 low versus NTF2 high dox+. Positive log_2_ fold changes indicate genes upregulated in NTF2 low. Negative log_2_ fold changes indicate genes downregulated in NTF2 low.

**Table S2: NTF2 high dox-versus NTF2 high dox+ transcriptomics data.** The RNAseq data are presented for NTF2 high dox-versus NTF2 high dox+. Positive log_2_ fold changes indicate genes upregulated in NTF2 high dox-. Negative log_2_ fold changes indicate genes downregulated in NTF2 high dox-.

**Table S3: VGP primary melanoma versus NTF2 low transcriptomics data.** The RNAseq data are presented for VGP primary melanoma versus NTF2 low. Positive log_2_ fold changes indicate genes upregulated in VGP primary melanoma. Negative log_2_ fold changes indicate genes downregulated in VGP primary melanoma.

## MOVIE LEGENDS

**Movie 1: NTF2 low cells exhibit fast cell migration.** A pipet tip was used to create a scratch through a completely confluent monolayer of NTF2 low cells. Bright-field imaging was performed over twelve hours.

**Movie 2: NTF2 high dox+ cells exhibit slow cell migration.** A pipet tip was used to create a scratch through a completely confluent monolayer of NTF2 high dox+ cells. Bright-field imaging was performed over twelve hours.

